# Kidney and Lung ACE2 expression after an ACE inhibitor or an Ang II receptor blocker: implications for COVID-19

**DOI:** 10.1101/2020.05.20.106658

**Authors:** Jan Wysocki, Enrique Lores, Minghao Ye, Maria Jose Soler, Daniel Batlle

## Abstract

**Background:** There have been concerns that ACE inhibitors and Ang II receptor blockers may cause an increase in ACE2, the main receptor for SARs-CoV-2.

**Methods:** Kidneys from two genetic models of kidney ACE ablation and mice treated with captopril or telmisartan were used to examine ACE2 in isolated kidney and lung membranes.

**Results:** In a global ACE KO mice, ACE2 protein abundance in kidney membranes was reduced to 42 % of wild type, p < 0.05. In ACE 8/8 mice that over-expresses cardiac ACE protein but also has no kidney ACE expression, ACE2 protein in kidney membranes was also decreased (38 % of the WT, p<0.01). In kidney membranes from mice that received captopril or telmisartan for 2 weeks there was a reduction in ACE2 protein (37% in captopril treated p<0.01) and 76% in telmisartan treated p <0.05). In lung membranes the expression of ACE2 was very low and not detected by western blotting but no significant differences in terms of ACE2 activity could be detected in mice treated with captopril (118% of control) or telmisartan (93% of control).

**Conclusions:** Genetic kidney ACE protein deficiency, suppressed enzymatic activity by Captopril or blockade of the AT1 receptor with telmisartan are all associated with a decrease in ACE2 in kidney membranes. ACE2 protein in kidney or lungs is decreased or unaffected by RAS blockers indicating that these medications can not pose a risk for SARS-CoV-2 infection related to amplification of ACE2 at these two target sites for viral entry.

## INTRODUCTION

Angiotensin I converting enzyme (ACE) is a dicarboxypeptidase that cleaves Angiotensin I (ANG I) to form angiotensin (ANG) II. Its enzymatically active homologue, Angiotensin Converting Enzyme 2 (ACE2), by contrast, degrades ANG II to form ANG-(1-7) as well as other substrates(1–5). There is limited information on whether organ-specific ACE2 expression is altered when ACE is decreased in genetic models where ACE is absent. If ACE2 acts indeed as a negative regulator of the renin-angiotensin system (RAS) by regulating angiotensin II levels then ACE2 should decrease when ACE decreases and conversely ACE2 activity should increase when ACE activity increases. This mode of physiologic regulation, however, can be altered in pathophysiologic conditions. For instance, it is known that in kidneys from diabetic mice ACE2 and ACE change in the opposite direction(6–9), such that in renal cortex from diabetic db/db (6, 7) and STZ mice(6) ACE is decreased, whereas ACE2 is increased. Of note, within the glomeruli from db/db mice(9) ACE and ACE2 are also altered in the opposite manner (high ACE and low ACE2) which is just the contrary of that seen in proximal tubules(6, 9). This finding in diabetic mice was generally confirmed in kidney biopsies from patients with diabetic kidney disease(10). To examine whether deficiency of kidney ACE results in the expected physiologic decrease in kidney ACE2 expression we used two different genetic models of ACE ablation. Captopril and telmisartan moreover were used to pharmacologically inhibit ACE activity and block the AT1 receptor, respectively.

The recent recognition that ACE2 is the main receptor that facilitates coronavirus entry into cells (11) raised concerns that treatment with renin-angiotensin system blockers (RAS) might increase the risk of developing a severe and fatal severe acute respiratory syndrome coronavirus-2 infection (12). This is relevant because certain groups of patients including those with hypertension, heart disease, diabetes mellitus, and clearly the elderly are at risk of COVID-19 and many of those patients are treated with RAS blockers (12–14). Concerns came from previous studies showing that angiotensin II type 1 receptor blockers and ACE inhibitors can upregulate ACE2 in certain experimental conditions (15–18). The evidence in this regard is not always consistent and differs among the diverse RAS blockers, animal species and differing organs.(19–24).

We reasoned that this issue should be investigated by assessing ACE2 expression using membrane membranes prepared from target organs where this protein resides. In polarized epithelia like the lungs, kidneys and intestine, ACE2 in its full length form is anchored to the apical plasma membrane(9). The kidney has abundant ACE2 in the apical border of the proximal tubule where it colocalizes with ACE(9). The lungs, by contrast, have a low level of ACE2 expression such that the metabolism of angiotensin II and formation of angiotensin 1-7 depends on enzymes other than ACE2, namely POP(25). To determine whether deficiency of kidney ACE alters kidney ACE2 expression we used two different genetic models of ACE ablation. To mimic the clinical setting of patients receiving RAS blockers are exposed to the current COVID-19 pandemic, captopril and telmisartan were administered for 2 weeks to pharmacologically inhibit ACE activity and the AT1 receptor, respectively.

## METHODS

### Animal Models

All studies were conducted with the review and approval of the Institutional Animal Care and Use Committee of Northwestern University. Organs from two genetic models of ACE kidney ablation (ACE.4 and ACE 8/8) were generously provided by Drs Hong D. Xiao and Kenneth Bernstein. ACE.4 mice have the somatic ACE promoter replaced by the kidney androgen-regulated protein promoter which in these mice is essentially non-functional(26); in the absence of exogenous androgens the levels of kidney ACE are less than 1% of normal and no ACE is detected in other organs; this model is overtly hypotensive (26). The effect of localized kidney ACE deficiency on kidney ACE2 levels was examined in ACE8/8 mouse which is a model lacking ACE in the kidney or vascular endothelium, but with 100-fold normal cardiac ACE levels(27). In addition, ACE8/8 mice have significant levels of ACE activity in the lung and in the plasma and blood (27). In this animal model, kidney represents a major change from wild-type mice as this organ normally expresses a substantial amount of ACE activity in both vascular endothelium and proximal tubular epithelium(27). The ACE8/8 mice have a near normal blood pressure and do not exhibit any gross abnormalities in kidney function(27). Both ACE models used in this study have a mixed C57/129 background(26, 27).

To examine the effect of pharmacological ACE inhibition on ACE2, two groups of 12-14 weeks old C57BLKS/J mice were assigned to drink either tap water (n=8) or tap water with an ACE inhibitor, captopril, (n=8) at a dose of 120 mg*kg^−1^*day^−1^ for 14 days. Two other experimental groups of 12-14 weeks old C57BLKS/J mice were randomly assigned to drink either tap water (vehicle, n=6) or tap water with an angiotensin II receptor antagonist, telmisartan (Boehringer Ingelheim), at a dose of 2 mg*kg^−1^*day^−1^ (n=6) for 14 days. Before captopril and telmisartan administration, mice were weighted and the daily fluid intake per mouse was recorded to estimate the concentration of the compound needed to be added to the drinking water. During the drug administration, water consumption and body weights were also controlled to make appropriate adjustments.

### RNA isolation and reverse transcriptase real-time PCR

RNA was isolated from kidney cortex with Trizol reagent (GIBCO Invitrogen). Quantitative real-time PCR was performed using the TaqMan Gold RT-PCR kit and ABI Prism 7700 (Applied Biosystems) sequence-detection system. Primers and probes for ACE were designed using Primer Express software (Applied Biosystems). The sequences of forward, reverse primer, and probe for ACE, ACE2 and Glyceraldehyde-3-phosphate dehydrogenase (G3PDH) were the same as described before(8). Reverse transcription was carried out for 30 min at 48°C. The ACE and ACE2 mRNA levels of the samples were normalized to their G3PDH contents. Experiments were carried out in triplicate for each data point.

### Protein extraction and measurement of enzymatic activity for ACE2

For total cell lysates, tissues (kidney cortex and lungs) were homogenized in a buffer consisting of (in mmol/l) 50 HEPES, pH 7.4, 150 NaCl, 0.5% Triton X-100, 0.025 ZnCl_2_, and 1.0 phenylmethylsulphonyl fluoride and then clarified by centrifugation at 6,000g for 15 min. For membrane fraction isolation, a previously employed protocol was used(7) but without EDTA addition. After measuring protein concentration, tissue samples were diluted in a buffer (50 mmol/l 4-morpholineethanesulfonic acid, 300 mmol/l NaCl, 10 μmol/l ZnCl_2_, and 0.01% Triton-X-100, pH 6.5), containing EDTA-free tablets. The plates were read using a fluorescence plate reader FLX800 (BIOTEK Instruments) at an excitation wavelength of 320 nm and an emission wavelength of 400 nm. All reactions were performed at ambient temperature in microtiter plates with a 100 μl total volume (6, 28).

### Western blot analysis

Total cell lysates and isolated membrane preparations from kidney cortices and lungs were isolated as specified above and subjected to Western blot analysis as previously described(6). For detection of ACE, nitrocellulose membranes were incubated with mouse monoclonal antibody (5C4, kind gift from Dr. Sergei Danilov). ACE2 protein was detected using our affinity purified rabbit anti-ACE2 antibody (obtained as described in. Signals on Western blots were quantified by densitometry (Eagle Eye, Stratagene) and corrected for GAPDH (total cell lysates) or β-actin (membrane fraction) loading control.

### Statistical analyses

The data were expressed as relative to the control (vehicle or wild-type) animal groups, assigning a value of 100% to the control baseline mean. The data were reported as mean ± standard error (SE). Significance was defined as p*< 0.05.* For comparisons between two independent means, the t-test was used.

## RESULTS

### ACE2 in kidneys from ACE.4 mice, a model of global ACE deficiency

In ACE.4 mice, kidney ACE mRNA was about 7±4% of the mRNA levels in WT mice as previously reported (26). Total cell lysates from ACE.4 mice kidney cortex were examined for the presence of ACE and ACE2 protein (Figure 1). The signal for ACE protein in kidneys from WT mice was detectable as a single band at the expected molecular weight of about 170 kD. In ACE.4 mice, ACE protein was barely detectable (equivalent of 3±1% of the WT) (Figure 1B). In ACE.4 mice, ACE2 protein abundance in kidney cortex total cell lysates and in isolated membranes was reduced to 70 % and to 42% of the WT, p<0.05, respectively (Figure 1B and 1C).

**Figure 1.**
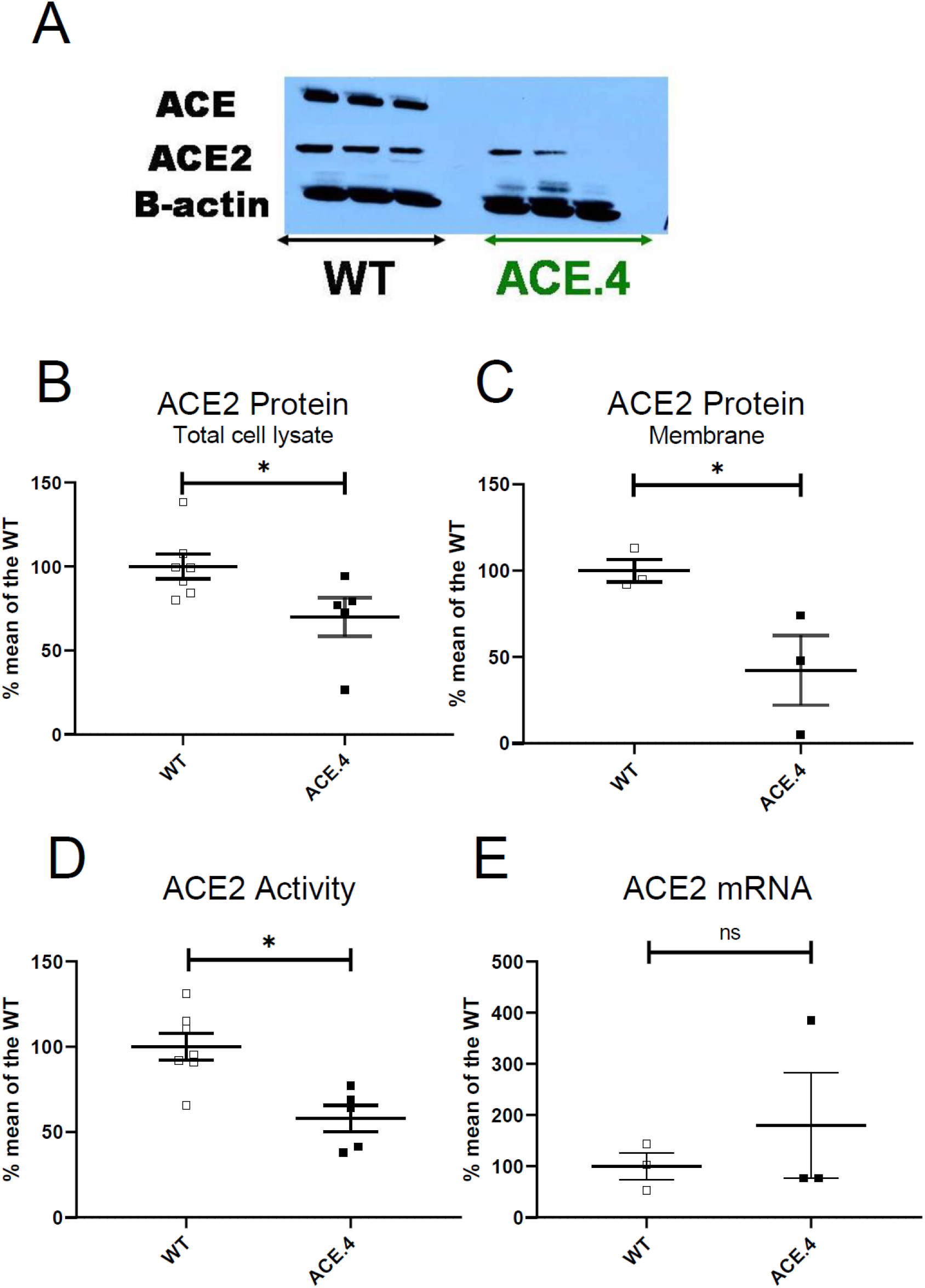
ACE2 protein, enzymatic activity and mRNA in kidneys from ACE.4 mice and wild type controls. Panel A shows a representative Western blot for ACE, ACE2 and β-actin for three mice from each respective group. ACE2 protein in kidney total cell lysates (panel B), ACE2 protein in isolated membranes (panel C), ACE2 enzymatic activity (panel D) and ACE2 mRNA (panel E) * p<0.05.).

Concordant with ACE2 protein expression, ACE2 activity was also reduced significantly in kidneys from ACE.4 mice as compared to WT controls (58.3±12% of the WT) (Figure 1D). By contrast, kidney cortex ACE2 mRNA levels in ACE.4 mice were not significantly different from those of their respective WT littermates (1.24±0.71 vs. 0.69±0.18 gene expression units, respectively) (Figure 1E).

### ACE2 in kidneys from ACE8/8 mice, a model of kidney ACE deficiency and cardiac ACE over-expression

Consistent with results from Xiao et al.(27), in kidney cortex from ACE8/8 mice there was no detectable ACE protein expression (Figure 2A), whereas it was clearly present in WT kidneys (Figure 2A). ACE2 protein expression in isolated kidney membranes from ACE8/8 was significantly reduced (38±9% of WT mice) (Figure 2B). In ACE8/8 mice, ACE2 activity likewise was lower than in the wild-type but this difference was small and not statistically significant (90±17% of the WT) (Figure 2C). ACE2 mRNA levels in kidney cortex from ACE8/8 mice were not different from those of their respective WT littermates (1.02±0.04 vs. 1.00±0.07, respectively).

**Figure 2.**
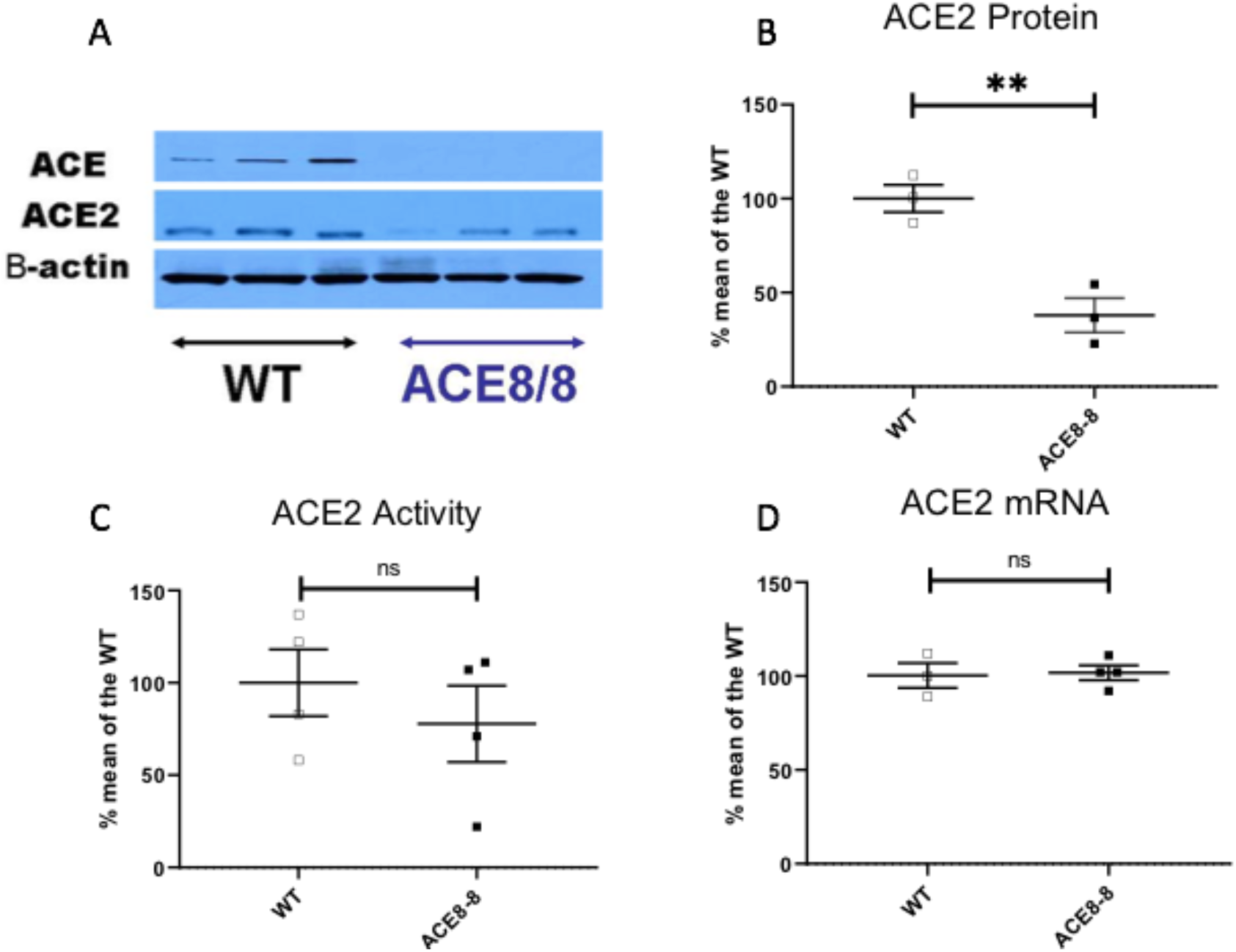
ACE2 protein, activity and m RNA in kidneys from ACE8-8 mice and wild type controls. Panel A shows a representative Western blot for ACE, ACE2 and β-actin for three mice from each respective group. Panel B shows ACE2 protein in isolated membranes. Panel C shows ACE2 enzymatic activity level and panel D ACE2 mRNA in cell lysates ** p<0.01.

### ACE2 in Kidneys from captopril and telmisartan treated mice

kidney cortex of mice treated with captopril or telmisartan for 14 days and respective vehicle treated controls were examined for the presence of ACE2 mRNA, protein and activity. No significant changes in mRNA levels were found between captopril treated and control mice (99±21%) of control. ACE2 activity was lower in isolated membrane from captopril-treated mice (81±8% of the vehicle-treated mice) but the difference did not reach statistical significance, (p = 0.053). ACE2 protein abundance in total lysates from captopril-treated mice was reduced to 71±5 of vehicle-treated mice but the difference did not reach statistical significance (Figure 3A). In isolated membranes, however, the decrease in ACE2 protein was profound and statistically significant (37±4% of the vehicle-treated mice) (Figure 3B). A corresponding increase in cytosolic ACE2 protein was found (Figure 3C). By confocal microscopy ACE2 is localized in the apical membrane where it strongly colocalizes with ACE as previously described site(9). A weak ACE2 staining can be seen in the cytoplasm of cells from a captopril treated mice suggesting internalization of the protein whereas ACE staining remains restricted to the luminal membrane (Figure 3C).

**Figure 3.**
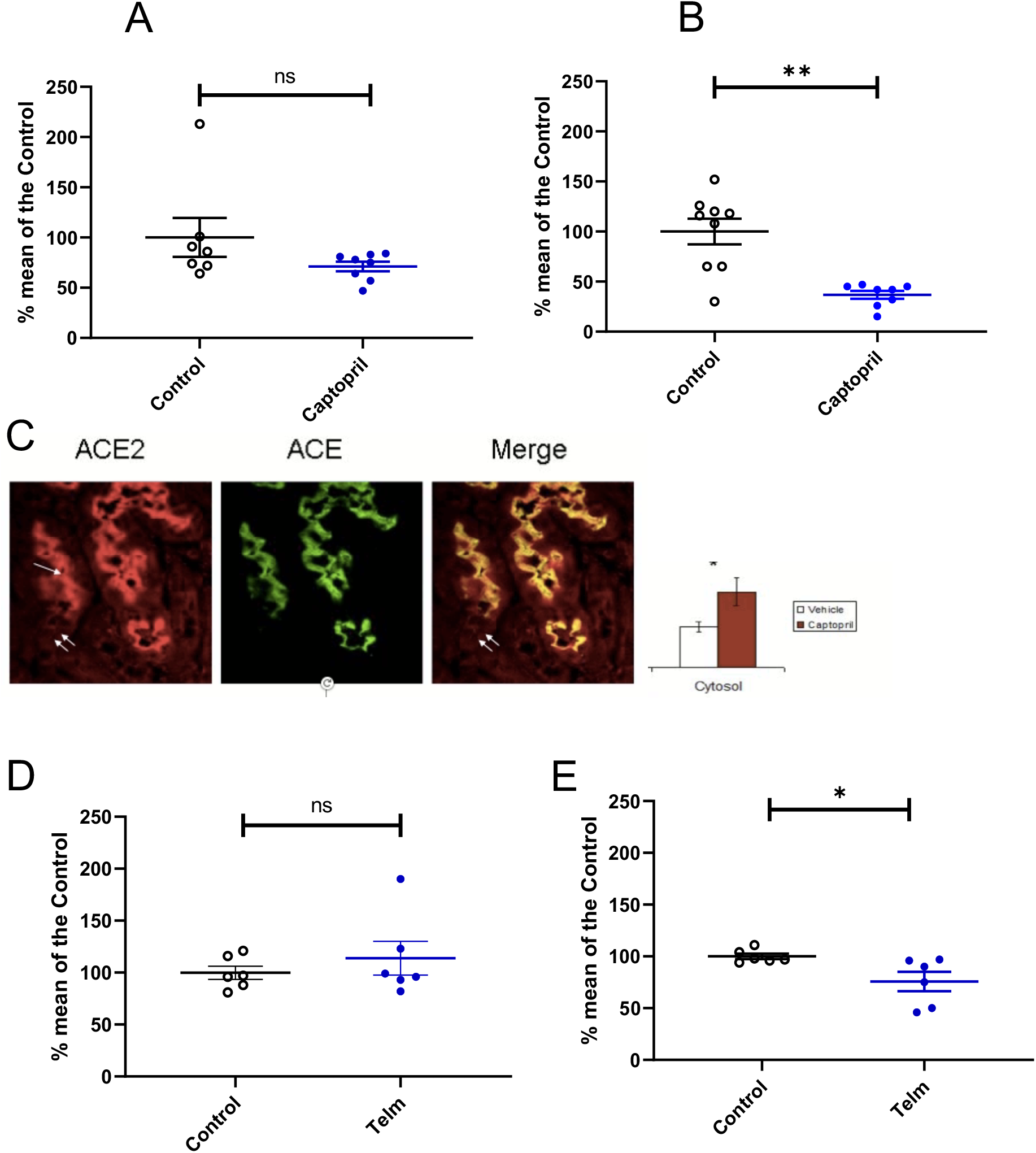
ACE2 protein expression in kidney cortex in kidneys from captopril and telmisartan treated mice. ACE2 protein in total lysates (Panel A) ACE2 protein in isolated membranes. (Panel B) Panel C. Proximal tubule confocal microscopy showing ACE2 (red) ACE (green) and merged image (yellow) ACE2 is the apical site (arrow), but some ACE2 staining is also seen in the cytoplasm (double arrow) of a captopril treated mice. The inset shows an increase in cytosolic ACE 2 protein in captopril treated as compared to vehicle treated mice. Panel D shows ACE2 protein in total cell lysates from telmisartan-treated mice and panel E shows ACE2 protein in isolated membranes from telmisartan-treated mice. * p<0.05, ** p<0.01.

ACE2 protein in lysates from telmisartan-treated mice was not significantly different from vehicle-treated mice (114±16%)(Figure 3D) whereas in isolated membranes there was a significant decrease in ACE2 protein (76± 9% of the vehicle-treated mice p<0.05) (Figure 3E).

### ACE2 in lungs from captopril and telmisartan treated mice

In lung tissue ACE2 protein is low as demonstrated by us recently(25) Attempts to perform western blots with either total lysates or isolated membranes did not produce a signal. Therefore, the results are limited to ACE2 activity which although low was consistently detected(figure 4). Captopril treatment had no significant effect on ACE2 activity either lung lysates (Figure 4A) or isolated membranes ( Figure 4B). Likewise, telmisartan had no significant effect on ACE2 activity in either lung lysates ( Figure 4C) or isolated membranes ( Figure 4D)

**Figure 4.**
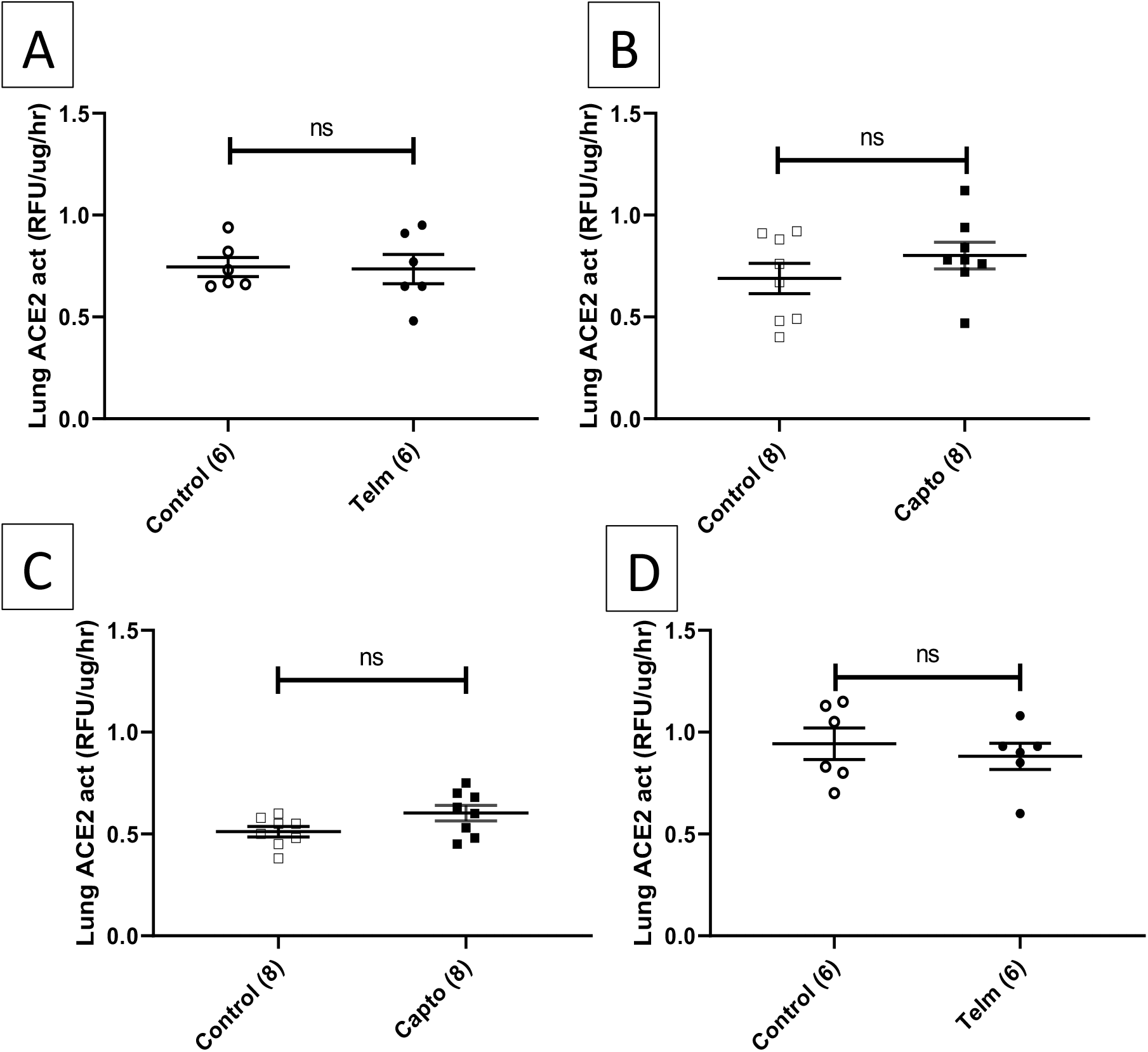
ACE2 activity in cell lysates (A and C) and isolated membrane preparations (B and D) from lungs of captopril and telmisartan treated mice. No significant differences were detected.

## DISCUSSION

Our findings in two different models of ACE genetic ablation, global in the ACE.4 mice and restricted to the kidney in the ACE8/8 mice shows that lack of ACE protein is associated with a significant reduction of kidney ACE2 protein. The decrease in ACE2 at the level of protein was not accompanied by a reduction in kidney ACE2 mRNA. This confirms that ACE2 can be regulated post transcriptionally as previously reported by us(6) and others(29). Our finding that ACE2 mRNA was unchanged in ACE deficient mice is also consistent with a previous report in another genetic model, the *tis*ACE-/-knockout mice, that lacks tissue ACE and possess about 40% of normal circulating levels of ACE(30). Similar to our results in two models of kidney ACE deficiency, in *tis*ACE-/-mice kidney expression of ACE2 mRNA was not altered by ACE depletion. ACE2 protein abundance or activity, however, were not determined in this previous study(30)

Altogether, these findings suggest that changes in ACE 2 occur post transcriptionally and are accompanied by changes in ACE2 in the same direction such a decrease in kidney ACE is apt to be associated with a decrease in kidney ACE2. Overall the directionally observed changes are consistent with a physiologic role of ACE2 in terms of regulating its main substrate, Angiotensin II (5). Given the intense interest on ACE2 as the SARS-CoV-2 receptor we next wanted to see if a similar directional change in ACE2 occurs when an ACE inhibitor or a AT1 blocker is chronically administered.

We reasoned that in terms of assessing ACE2 as the SARS-CoV-2 receptor what matters the most is the relative abundance of full-length membrane bound ACE2 protein. In polarized epithelia like the lungs, kidneys and intestine, ACE2 in its full length form is anchored to the apical plasma membrane(9). The effect of ACEi and ARBs on ACE2 protein expression in the lungs has not been previously studied and previous studies in kidney and other organs gave variable results(15–23). Here we show that captopril and telmisartan both decrease kidney ACE2 protein in isolated membranes without significantly affecting protein abundance in total cell lysates (Figure 3). Captopril in particular produced a marked decline in ACE2 protein in isolated membranes while increasing cytosolic ACE2(Figure 3C). The latter result was somewhat unexpected because Ang II has been shown to produce internalization of ACE2 in cultured cells (31). The discrepancy between this previous study in ACE2-transfected neuroblastoma cells and our findings with in vivo chronic administration of captopril are not apparent to us but clearly the experimental conditions are widely different.

In lung tissue, the expression of ACE protein is very low(25). Consistent with this previous study we could not detect ACE2 by western analysis in lung lysates or isolated membranes. In terms of enzymatic ACE2 activity there was no significant effect by either captopril or telmisartan. (Figure 4). Within lung tissue, ACE2 protein is detectable only in type II pneumocytes which makes it very difficult to assess the impact of RAS blockers using lung membranes.

In conclusion, genetic ablation and inhibition of ACE are accompanied by a reduction of ACE2 protein and enzymatic activity. In a transgenic model of cardiac ACE overexpression and kidney ACE ablation, ACE2 protein in the kidney was also decreased. This suggest that changes in the expression in one enzyme may elicit similarly directional changes in the other homologue such that formation and degradation of ANG II can be coupled. The administration of an ACE inhibitor and an AT1 receptor blocker decreased ACE2 protein expression in kidney isolated membranes and had no detectable effect on ACE2 activity in lung isolated membranes where ACE2 expression is very low. These findings altogether suggest that two potential target sites for SARS-CoV-2 infection where ACE2 is expressed, the kidney and lung apical membrane, RAS blockers decrease or had no effect on ACE2 expression. We conclude that these medications do not increase ACE2 expression in lung or kidney epithelia and therefore can no longer be considered to pose a risk for COVID-19. This information supports the position of many medical societies and recent publications expressing the view that RAS blockers should not be abandoned in COVID-19 patients or the large number of people that can be potentially exposed.

## References

1. Tipnis SR, Hooper NM, Hyde R, Karran E, Christie G, Turner AJ: A human homolog of angiotensin-converting enzyme. Cloning and functional expression as a captopril-insensitive carboxypeptidase. J Biol Chem, 275: 33238–33243, 2000

2. Donoghue M, Hsieh F, Baronas E, Godbout K, Gosselin M, Stagliano N, Donovan M, Woolf B, Robison K, Jeyaseelan R, Breitbart RE, Acton S: A novel angiotensin-converting enzyme-related carboxypeptidase (ACE2) converts angiotensin I to angiotensin 1-9. Circ Res, 87: E1–9, 2000

3. Ortiz-Melo DI, Gurley SB: Angiotensin converting enzyme 2 and the kidney. Current opinion in nephrology and hypertension, 25: 59–66, 2016

4. Li N, Zimpelmann J, Cheng K, Wilkins JA, Burns KD: The role of angiotensin converting enzyme 2 in the generation of angiotensin 1–7 by rat proximal tubules. American Journal of Physiology-Renal Physiology, 288: F353–F362, 2005

5. Batlle D, Wysocki J, Soler MJ, Ranganath K: Angiotensin-converting enzyme 2: enhancing the degradation of angiotensin II as a potential therapy for diabetic nephropathy. Kidney international, 81: 520–528, 2012

6. Wysocki J, Ye M, Soler MJ, Gurley SB, Xiao HD, Bernstein KE, Coffman TM, Chen S, Batlle D: ACE and ACE2 activity in diabetic mice. Diabetes, 55: 2132–2139, 2006

7. Ye M, Wysocki J, Naaz P, Salabat MR, LaPointe MS, Batlle D: Increased ACE 2 and decreased ACE protein in renal tubules from diabetic mice: a renoprotective combination? Hypertension, 43: 1120–1125, 2004

8. Soler M, Wysocki J, Ye M, Lloveras J, Kanwar Y, Batlle D: ACE2 inhibition worsens glomerular injury in association with increased ACE expression in streptozotocin-induced diabetic mice. Kidney international, 72: 614–623, 2007

9. Ye M, Wysocki J, William J, Soler MJ, Cokic I, Batlle D: Glomerular localization and expression of angiotensin-converting enzyme 2 and angiotensin-converting enzyme: implications for albuminuria in diabetes. Journal of the American Society of Nephrology, 17: 3067–3075, 2006

10. Mizuiri S, Hemmi H, Arita M, Ohashi Y, Tanaka Y, Miyagi M, Sakai K, Ishikawa Y, Shibuya K, Hase H, Aikawa A: Expression of ACE and ACE2 in individuals with diabetic kidney disease and healthy controls. American journal of kidney diseases : the official journal of the National Kidney Foundation, 51: 613–623, 2008

11. Hoffmann M, Kleine-Weber H, Schroeder S, Krüger N, Herrler T, Erichsen S, Schiergens TS, Herrler G, Wu N-H, Nitsche A: SARS-CoV-2 cell entry depends on ACE2 and TMPRSS2 and is blocked by a clinically proven protease inhibitor. Cell, 2020

12. Zheng Y-Y, Ma Y-T, Zhang J-Y, Xie X: COVID-19 and the cardiovascular system. Nature Reviews Cardiology, 17: 259–260, 2020

13. Huang C, Wang Y, Li X, Ren L, Zhao J, Hu Y, Zhang L, Fan G, Xu J, Gu X: Clinical features of patients infected with 2019 novel coronavirus in Wuhan, China. The lancet, 395: 497–506, 2020

14. Fang L, Karakiulakis G, Roth M: Are patients with hypertension and diabetes mellitus at increased risk for COVID-19 infection? The Lancet Respiratory Medicine, 2020

15. Wang X, Ye Y, Gong H, Wu J, Yuan J, Wang S, Yin P, Ding Z, Kang L, Jiang Q: The effects of different angiotensin II type 1 receptor blockers on the regulation of the ACE-AngII-AT1 and ACE2-Ang (1–7)-Mas axes in pressure overload-induced cardiac remodeling in male mice. Journal of molecular and cellular cardiology, 97: 180–190, 2016

16. Ferrario CM, Jessup J, Chappell MC, Averill DB, Brosnihan KB, Tallant EA, Diz DI, Gallagher PE: Effect of angiotensin-converting enzyme inhibition and angiotensin II receptor blockers on cardiac angiotensin-converting enzyme 2. Circulation, 111: 2605–2610, 2005

17. Soler MJ, Ye M, Wysocki J, William J, Lloveras J, Batlle D: Localization of ACE2 in the renal vasculature: amplification by angiotensin II type 1 receptor blockade using telmisartan. American Journal of Physiology-Renal Physiology, 296: F398–F405, 2009

18. Soler MJ, Barrios C, Oliva R, Batlle D: Pharmacologic modulation of ACE2 expression. Current hypertension reports, 10: 410, 2008

19. Mancia G, Rea F, Ludergnani M, Apolone G, Corrao G: Renin–angiotensin–aldosterone system blockers and the risk of Covid-19. New England Journal of Medicine, 2020

20. Danser AJ, Epstein M, Batlle D: Renin-angiotensin system blockers and the COVID-19 pandemic: at present there is no evidence to abandon renin-angiotensin system blockers. Hypertension : HYPERTENSIONAHA. 120.15082, 2020

21. Vaduganathan M, Vardeny O, Michel T, McMurray JJ, Pfeffer MA, Solomon SD: Renin–angiotensin– aldosterone system inhibitors in patients with Covid-19. New England Journal of Medicine, 382: 1653–1659, 2020

22. Lely A, Hamming I, van Goor H, Navis G: Renal ACE2 expression in human kidney disease. The Journal of Pathology: A Journal of the Pathological Society of Great Britain and Ireland, 204: 587–593, 2004

23. South AM, Diz DI, Chappell MC: COVID-19, ACE2, and the cardiovascular consequences. American Journal of Physiology-Heart and Circulatory Physiology, 318: H1084–H1090, 2020

24. Perico L, Benigni A, Remuzzi G: Should COVID-19 concern nephrologists? Why and to what extent? The emerging impasse of angiotensin blockade. Nephron, 144: 213–221, 2020

25. Serfozo P, Wysocki J, Gulua G, Schulze A, Ye M, Liu P, Jin J, Bader M, Myöhänen T, García-Horsman JA: Ang II (angiotensin II) conversion to angiotensin-(1-7) in the circulation is POP (prolyloligopeptidase)-dependent and ACE2 (angiotensin-converting enzyme 2)-independent. Hypertension, 75: 173–182, 2020

26. Campbell DJ, Alexiou T, Xiao HD, Fuchs S, McKinley MJ, Corvol P, Bernstein KE: Effect of reduced angiotensin-converting enzyme gene expression and angiotensin-converting enzyme inhibition on angiotensin and bradykinin peptide levels in mice. Hypertension, 43: 854–859, 2004

27. Xiao HD, Fuchs S, Campbell DJ, Lewis W, Dudley Jr SC, Kasi VS, Hoit BD, Keshelava G, Zhao H, Capecchi MR: Mice with cardiac-restricted angiotensin-converting enzyme (ACE) have atrial enlargement, cardiac arrhythmia, and sudden death. The American journal of pathology, 165: 1019–1032, 2004

28. Ye M, Wysocki J, Gonzalez-Pacheco FR, Salem M, Evora K, Garcia-Halpin L, Poglitsch M, Schuster M, Batlle D: Murine recombinant angiotensin-converting enzyme 2: effect on angiotensin II-dependent hypertension and distinctive angiotensin-converting enzyme 2 inhibitor characteristics on rodent and human angiotensin-converting enzyme 2. Hypertension, 60: 730–740, 2012

29. Lambert DW, Lambert LA, Clarke NE, Hooper NM, Porter KE, Turner AJ: Angiotensin-converting enzyme 2 is subject to post-transcriptional regulation by miR-421. Clin Sci (Lond), 127: 243–249, 2014

30. Modrall JG, Sadjadi J, Brosnihan KB, Gallagher PE, Yu C-h, Kramer GL, Bernstein KE, Chappell MC: Depletion of tissue angiotensin-converting enzyme differentially influences the intrarenal and urinary expression of angiotensin peptides. Hypertension, 43: 849–853, 2004

31. Deshotels MR, Xia H, Sriramula S, Lazartigues E, Filipeanu CM: Angiotensin II mediates angiotensin converting enzyme type 2 internalization and degradation through an angiotensin ii type i receptor–dependent mechanism. Hypertension, 64: 1368–1375, 2014

